# The basis of sharp spike onset in standard biophysical models

**DOI:** 10.1101/037549

**Authors:** Maria Telenczuk, Bertrand Fontaine, Romain Brette

**Affiliations:** Sorbonne Universités, UPMC Univ Paris 06, INSERM, CNRS, Institut de la Vision, 17 rue Moreau, 75012 Paris, France; Laboratory of Auditory Neurophysiology, University of Leuven, 3000 Leuven, Belgium

## Abstract

In most vertebrate neurons, spikes initiate in the axonal initial segment (AIS). When recorded in the soma, they have a surprisingly sharp onset, as if sodium (Na) channels opened abruptly. The main view stipulates that spikes initiate in a conventional manner at the distal end of the AIS, then progressively sharpen as they backpropagate to the soma. We examined the biophysical models used to substantiate this view, and we found that orthodromic spikes do no initiate through a local axonal current loop that propagates along the axon, but through a global current loop encompassing the AIS and soma, which forms an electrical dipole. Therefore, the phenomenon is not adequately modeled as the backpropagation of an electrical wave along the axon, since the wavelength would be as large as the entire system. Instead, in these models, we found that spike initiation rather follows the critical resistive coupling model proposed recently, where the Na current entering the AIS is matched by the axial resistive current flowing to the soma. Besides demonstrating it by examining the balance of currents at spike initiation, we show that the observed increase in spike sharpness along the axon is artifactual and disappears when an appropriate measure of rapidness is used; instead, somatic onset rapidness can be predicted from spike shape at initiation site. Finally, we reproduce the phenomenon in a two-compartment model, showing that it does not rely on propagation. In these models, the sharp onset of somatic spikes is therefore not an artifact of observing spikes at the incorrect location, but rather the signature that spikes are initiated through a global soma-AIS current loop forming an electrical dipole.

**Author summary:** In most vertebrate neurons, spikes are initiated in the axonal initial segment, next to the soma. When recorded at the soma, action potentials appear to suddenly rise as if all sodium channels opened at once. This has been previously attributed to the backpropagation of spikes from the initial segment to the soma. Here we demonstrate with biophysical models that backpropagation does not contribute to the sharpness of spike onset. Instead, we show that the phenomenon is due to the resistive coupling between the large somatodendritic compartment and the small axonal compartment, a geometrical discontinuity that leads to an abrupt variation in voltage.

## INTRODUCTION

In most vertebrate neurons, action potentials are generated by the opening of sodium (Na) channels in the axon initial segment (AIS) (Debanne et al., 2011). According to the standard textbook account, spikes initiate through the interplay between two local transmembrane currents, when the inward Na current exceeds the outward leak current, carried mostly by potassium (K) (Fig. 1A). Because macroscopically Na channels open gradually with depolarization (Boltzmann slope factor: k_a_ ≈ 6 mV (Kole et al., 2008)), spike onset appears smooth in standard isopotential neuron models (Fig. 1B, top left). In contrast, the onset of spikes recorded at the soma of cortical neurons appears very sharp: in a voltage trace, spikes appear to suddenly rise from resting potential (Naundorf et al., 2006) (Fig. 1B, bottom, human cortical pyramidal neuron from Testa-Silva et al., 2014), as if all Na channels opened at once.

**Figure 1.**
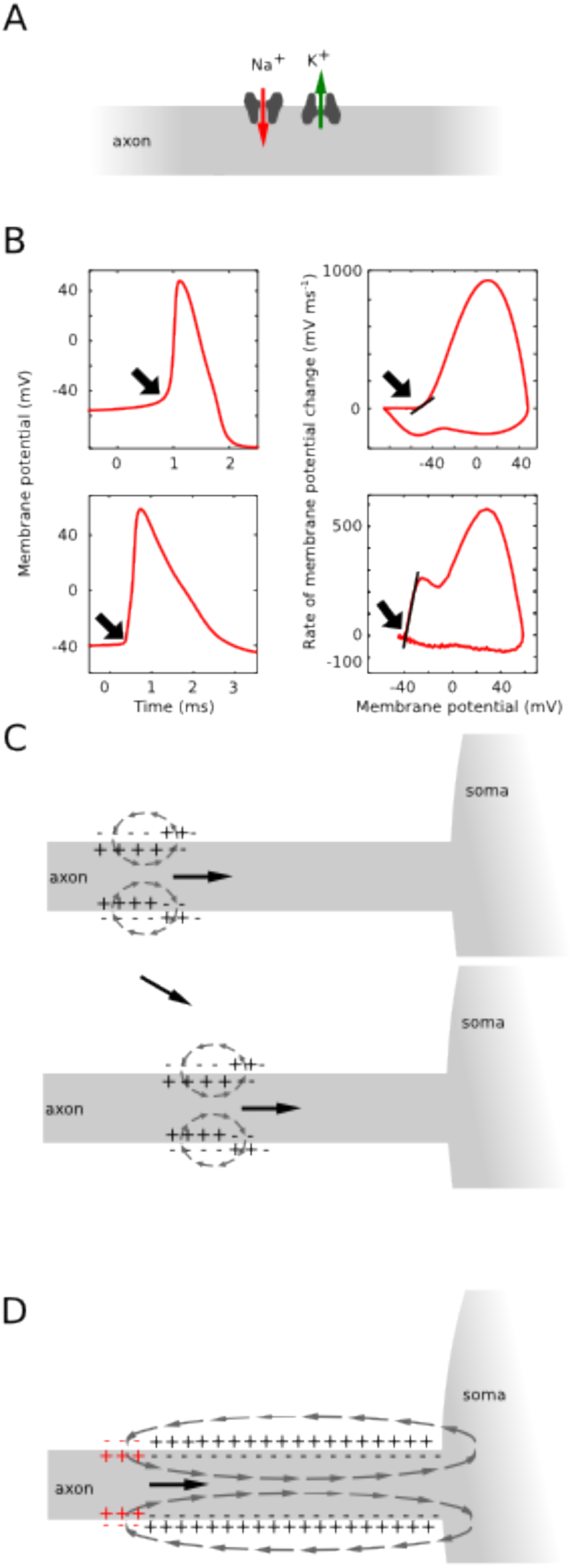
Theories of spike initiation. A, Standard account of spike initiation: spike initiation results from the interplay between Na current and K current (mostly leak) flowing through the membrane at the initiation site. B, Top: The isopotential Hodgkin-Huxley model produces spikes with smooth onset (left), exhibiting a gradual increase in dV/dt as a function of membrane potential V (right: onset rapidness measured as the slope at 20 mV/ms = 5.6 ms^-1^). Bottom: cortical neurons have somatic spikes with sharp onsets (left), with steep increase in dV/dt as a function of V (onset rapidness: 28.8 ms^-1^; human cortical data from Testa et al. (2014)). C, Backpropagation hypothesis: spikes are initiated according to the conventional account, with a local axonal current loop propagating towards the soma. D, Critical resistive coupling hypothesis: owing to the strong resistive coupling between the two sites and the soma acting as a current sink, spike initiation results from the interplay between Na current and axial current. Spikes then initiate through a global current loop encompassing AIS and soma, which behaves as an electrical dipole,

It has been proposed that Na channels in the AIS cooperate, so that they actually open all at once instead of gradually as a function of local voltage (Naundorf et al., 2006; Öz et al., 2015). However, this phenomenon has not been observed in the AIS (see Discussion). In addition, detailed multicompartmental models with standard biophysics can exhibit sharp somatic spikes (McCormick et al., 2007; Yu et al., 2008), when Na channel density is high enough (Baranauskas et al., 2010). According to the *backpropagation hypothesis,* this phenomenon is due to the progressive sharpening of spike onset between the axonal initiation site and the soma, partly due to the Na channels placed between the two sites (Fig. 1C; Yu et al., 2008). In this view, spike initiation follows the standard account: a spike initiates at a distal point of the AIS through a local current loop, and the electrical wave progressively propagates along the axon towards the soma, while its shape becomes sharper. In this view, the somatic kink does not bear any significance for excitability, since it only appears after spike initiation. This point is disputed because input-output properties and other features of excitability do not empirically match the predictions of standard accounts of spike initiation (Ilin et al., 2013; Brette, 2015).

A theoretical study proposed a different view, which we will call *critical resistive coupling* (Brette, 2013). The soma acts as a current sink for the initiation site because of the size difference and the short distance between the two sites. It follows that the Na current at spike initiation is not opposed by local transmembrane currents (the leak current), but by the resistive axial current flowing to the soma (Fig. 1D). Consequently, spikes initiate through a global current loop that encloses the AIS and soma, rather than a local axonal current loop: Na current through the AIS, resistive current between soma and AIS, capacitive current through the soma, and resistive current in the extracellular space to close the loop. Thus the soma and AIS form an electrical dipole at spike initiation, and it is therefore not appropriate to speak of wave propagation (the wavelength would be the entire system). When the product of axial resistance and Na conductance is greater than a critical value, Na channels open as a discontinuous function of somatic voltage, with consequences not only on somatic spike shape but also on input-output properties of neurons (Brette, 2015). This explanation attributes no role to active backpropagation or to the somatodendritic capacitance, beyond the requirement that the capacitance must be large enough for the soma to act as a current sink. Which of the critical resistive coupling and backpropagation hypotheses applies has not been determined in detailed biophysical models.

We therefore examined multicompartmental models of spike initiation, including models that were previously used to substantiate the backpropagation hypothesis, to determine which of the critical resistive coupling and backpropagation hypotheses actually applies. We found that: 1) the soma and AIS form an electrical dipole at spike initiation; 2) Na channels open as a discontinuous function of somatic voltage; 3) at spike initiation, the main current opposing the Na current is the axial resistive current; 4) excitability increases with intracellular resistivity; 5) active backpropagation is neither sufficient nor necessary for sharp spike initiation; 6) provided the somatodendritic compartment is large enough, its size has no quantitative impact on somatic onset rapidness; 7) the apparent sharpening of spikes as they backpropagate to the soma is an artifact of the measure of rapidness; in contrast, somatic onset rapidness can be predicted from spike shape at the initiation site. Finally, we show that the phenomenon can be reproduced by a model with only two resistively coupled compartments and standard channel properties. We conclude that in standard biophysical models, the biophysical basis of sharp somatic spikes is not backpropagation but critical resistive coupling, where spikes initiate through a global current loop encompassing the soma and AIS, rather than local current loops propagating towards the soma. This implies that in these models the sharpness of spike initiation is not an artifact, but a feature of normal (orthodromic) spike initiation through the soma-AIS dipole.

## RESULTS

### Intracellular and extracellular features of sharp spike initiation in multicompartmental models

We examined two multicompartmental Hodgkin-Huxley models that display somatic spikes with sharp onset (Yu et al., 2008; see Methods): one with an idealized morphology consisting of a uniform cylindrical axon (diameter 1 μm, length 50 μm) and a larger cylindrical soma (Fig. 2, left column), and one with the reconstructed morphology of a cortical pyramidal cell (Fig. 2, middle column). Action potentials are initiated in the axon, which has a high density of Na channels (8000 pSμm^2^), and regenerated in the soma, which has a lower density of Na channels (800 pSμm^2^). In both models, voltage traces show a distinct “kink” at the onset of somatic spikes (Fig. 2A, top, orange), which appears also when somatic Na channels are blocked (dotted orange). This kink is not present in the axonal spike (blue). Phase plots of dV/dt versus membrane potential V are biphasic (Fig. 2A, bottom), with a first component corresponding to the axonal Na current and a second component due to the somatic Na current. The sharpness of spike onset appears as a steep slope at threshold in the phase plots, called “initial phase slope” or “onset rapidness” (Na channels in the soma: 52.5 ms^-1^ and 52 ms^-1^ in the simple and detailed model, respectively; no Na channels in the soma: 55.6 ms^-1^ and 54 ms^-1^; compare with Fig. 1B: 5.6 ms^-1^; see Methods). We also examined spike initiation in the same way in another multicompartmental model of rat cortical pyramidal cell, where the models of Na and K currents were fitted to patch-clamp measurements in those neurons, separately in soma and AIS (Hallermann et al., 2012). The same qualitative features were observed (Fig. 2, right column).

**Figure 2.**
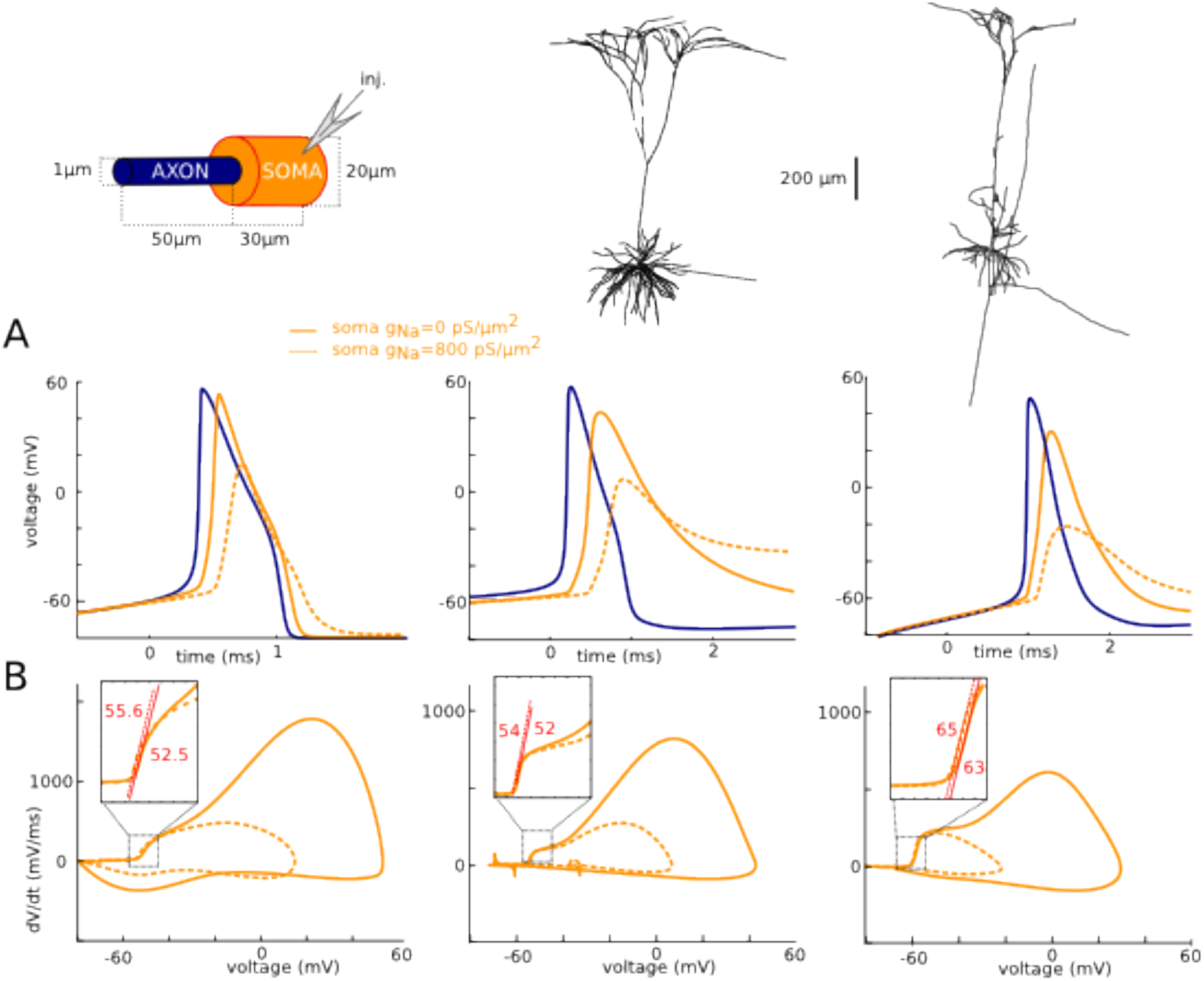
Intracellular features of sharp spike initiation in multicompartmental models. Left: model with simplified soma-axon geometry. Middle: cortical pyramidal cell model with morphological reconstruction from Yu et al. (2008). Right: pyramidal cell model from Hallermann et al. (2012). A, Somatic voltage trace of a spike, with (solid orange) and without (dotted orange) somatic Na channels, and axonal spike (blue). B, Phase plot of the same trace, showing dV/dt versus membrane potential V.

### Extracellular field at spike initiation

We then examined the extracellular field at spike initiation in the simple model (Fig. 3 and Movie S1). At the very beginning of the spike (Fig. 3A), the current injected at the soma is seen to exit the soma and part of it enters the distal end of the AIS. At the peak of the distal axonal spike (Fig. 3B), which corresponds to the knee of the somatic spike, the electrical field clearly shows that the soma and AIS form an electrical dipole, with current entering the AIS, flowing to the soma, exiting at the soma and returning through the extracellular space. During repolarization (Fig. 3C), the soma-AIS dipole is inverted, with current flowing intracellularly from soma to AIS, and extracellularly from AIS to soma.

**Figure 3.**
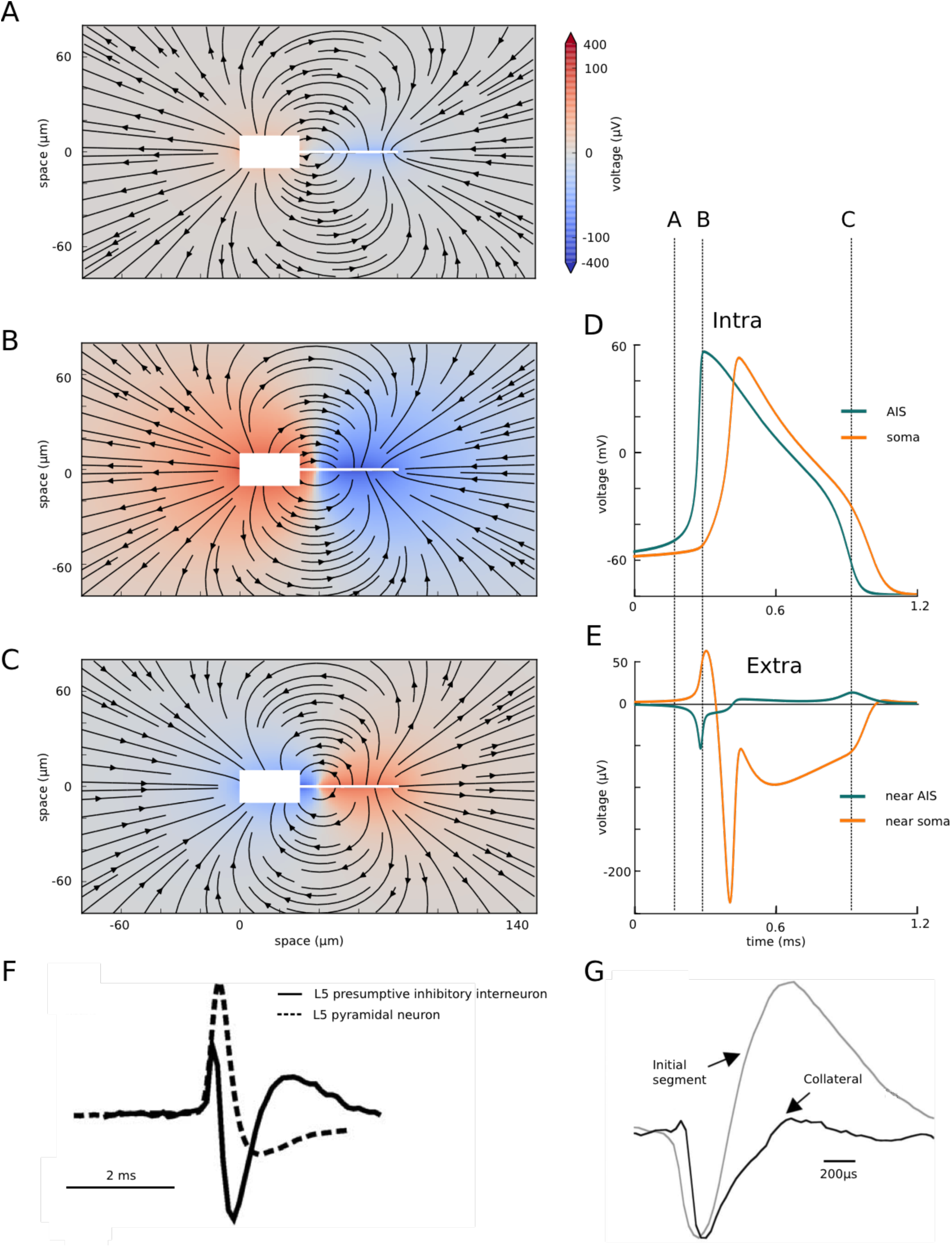
Extracellular field at spike initiation. A-C, Extracellular potential (color coded) and electrical field (arrows) around the simplified neuron (white box and line), at three different times indicated in D and E. D, Intracellular voltage trace at the soma and AIS distal end. E, Extracellular potential near the soma and AIS distal end. F, Extracellular recording near the soma of two cortical neurons (from Joshi and Hawken, 2006). G, Extracellular AP recording near the AIS (grey) of a cortical pyramidal cell (from Palmer and Stuart, 2006).

It is important to notice that even though the intracellular spike appears at the soma after a substantial delay following the axonal spike (Fig. 3D), the electrical dipole forms very quickly over the entire system (Fig. 3E). This discrepancy is due to the fact that the soma has a large capacitance and therefore a large charging time. Thus, the delay between the axonal and somatic spikes is better understood as a charging time (as an electrode charging a cell) rather than a propagation time.

The formation of the soma-AIS dipole at spike initiation manifests itself as negative extracellular potential near the AIS (current entering the AIS) and positive extracellular potential near the soma (current leaving the soma), as seen in Fig. 3E. Near the soma, this is followed by a negative peak corresponding to the somatic spike, and a smaller positive peak corresponding to the repolarization. Near the AIS, the negative peak is followed by a positive potential corresponding to the somatic spike (inversion of the dipole) and the axonal repolarization. Fig. 3F–G show extracellular recordings next to the soma and AIS, recorded experimentally in two different neurons. There are clearly quantitative differences with Fig. 3E, which depend both on the particular model and the position of the extracellular electrodes, but the same extracellular features of a dipole are seen. However, the precise temporal relationship between the extracellular waveforms of the two sites would be necessary to draw firm conclusions.

### Currents at spike initiation

The sharpness of spike initiation is not only seen in the initial shape of action potentials. It also appears in voltage-clamp (Fig. 4). In the same 3 models used in Fig. 2, we recorded currents in somatic voltage-clamp, with an ideal configuration (access resistance R_s_ = 0.1 MQ), varying the command potential from 1 mV below spike threshold to 5 mV above it (Fig. 3A). This type of recording can be challenging in practice because the pipette resistance introduces artifacts when passing large currents (see Discussion), but this was not the case in these simulations. We notice that recorded currents do not increase gradually with voltage, as when recording an isopotential patch of membrane, but instead have all-or-none characteristics: there is no current 1 mV below threshold (black), and a very large current (10-20 nA) just above threshold (red). Current amplitude varies little, but latency decreases when command potential is increased. We can see that this current mirrors the membrane potential in the distal AIS, where a spike is produced (Fig. 4A, bottom). Thus these currents recorded in somatic voltage-clamp correspond to axial currents coming from the AIS.

Plotting the peak current as a function of command potential shows that the peak current increases discontinuously when the voltage command exceeds a threshold value (Fig. 4B, top), which is close to the voltage at spike onset measured in current-clamp (-60 mV, -58.2 mV and - 62.2 mV in voltage-clamp vs. -59.7 mV, -56 mV and -65.3 mV in current-clamp, when dV/dt = 5 mV/ms). Blocking the somatic Na channels has no effect on this discontinuity (dotted orange). This discontinuity in the current-voltage relationship corresponds to a discontinuity in the proportion of open Na channels in the initiation site (distal end of the axon) as a function of somatic voltage (Fig. 4B, bottom), even though the activation curve of Na channels is smooth: effectively, Na channels open simultaneously as a function of somatic (but not axonal) voltage.

At first sight, it might seem trivial that a spike is produced when the somatic voltage exceeds a threshold. Yet, this is not the case in case in isopotential models, spatially extended models with somatic initiation (Koch et al., 1995), or axonal models (Jack et al., 1975), where a charge or a voltage threshold may apply depending on cases. A voltage threshold is a defining feature of integrate-and-fire models and a property of models with distal axonal initiation (Brette, 2015), but only when the axial resistance between soma and axonal initiation site is large enough (hence the name critical resistive coupling) (Brette, 2013). Therefore, the phenomenon of sharp initiation is not only about spike shape, but also about the way the axonal current changes with somatic voltage. Specifically, a very small change in somatic voltage can produce a very large change in axonal current. This phenomenon is the basic feature of the critical resistive coupling model (Brette, 2013), which also predicts the inverse relation between latency and distance to threshold characteristic of a saddle-node bifurcation. It can be reproduced by a two-compartment model (Milescu et al., 2010, Fig. 3A).

Empirically, this phenomenon has been observed in motoneurons (Barrett and Crill, 1980), cortical pyramidal cells and inferior olivary neurons (Milescu et al., 2010). An example is shown on Fig. 4C in a raphé neuron (from Milescu et al., 2010). Peak current shows a discontinuity when command potential is increased (right, red), except when axonal Na channels are inactivated with a prepulse (black).

We then examined the balance of currents near threshold at the axonal initiation site (Fig. 5). In all three models, the main current opposing the Na current (red) is the axial current flowing to the soma (black), while the K current (green) becomes significant only near the peak of the axonal spike. In this figure, transmembrane currents are summed over the AIS membrane, and the axial current is measured at the soma-axon junction. Thus, at spike initiation, the initial dynamics of the spike is determined by the interaction between the axial and Na current, rather than between the K (mainly leak) and Na current. This reflects the fact that current flows intracellularly between the two poles of the soma-AIS dipole (Fig. 3). If follows in particular that axial resistance rather than membrane resistance should determine excitability.

**Figure 4.**
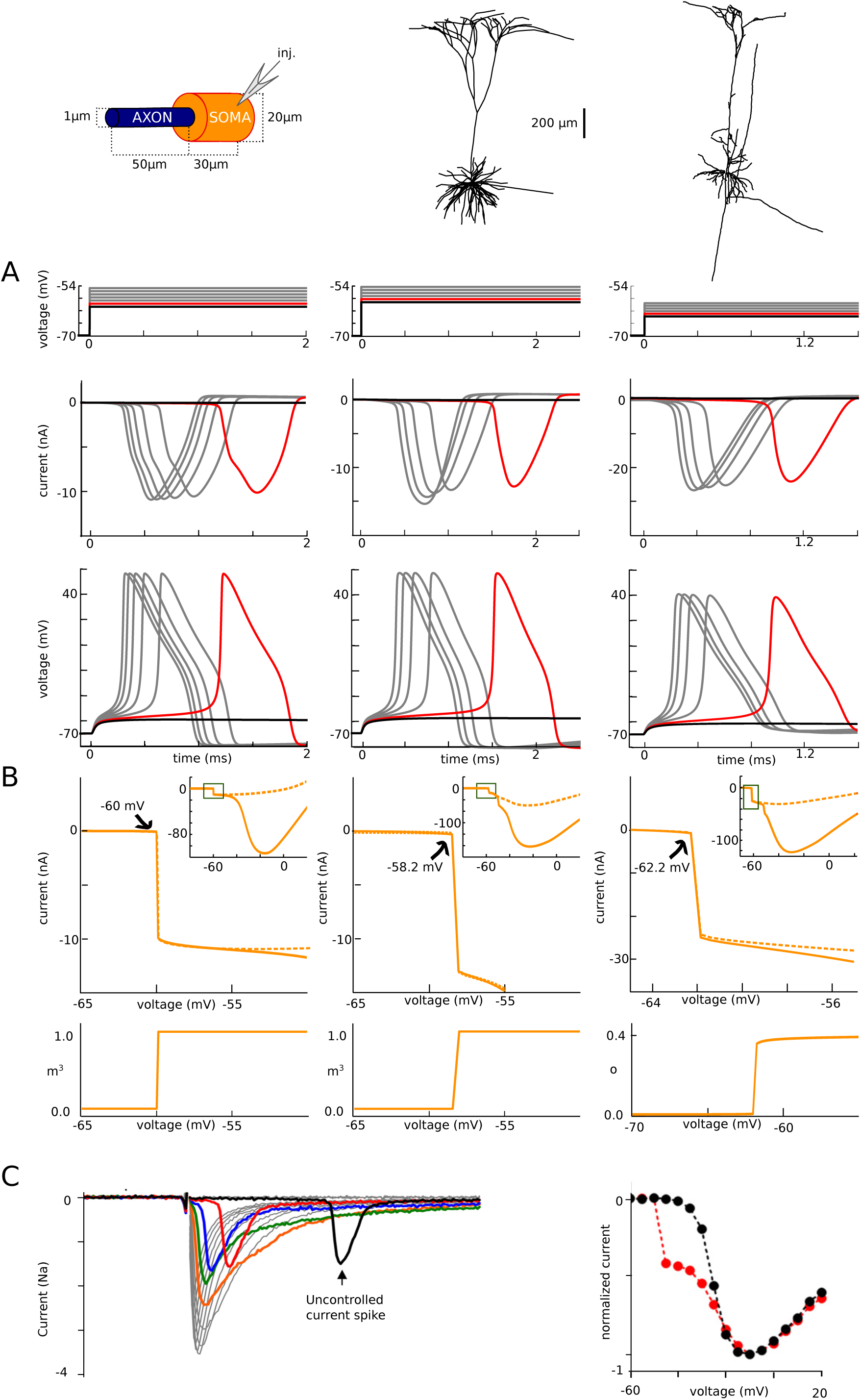
Currents at spike initiation. A, Somatic voltage-clamp recordings. Top: somatic membrane potential, spaced by 1 mV increments from threshold (red), with one trace just below threshold. Middle: recorded currents. Bottom: membrane potential at the AIS end. B, Top: peak current measured in somatic voltage-clamp versus holding voltage, with and without somatic Na channels, showing a discontinuity. Bottom: peak proportion of open Na channels at the distal axonal end versus holding voltage (variable m^3^ representing activation is shown for the first two models; variable o representing current-passing state is shown for the third model). C, Left, Current traces experimentally measured in somatic voltage-clamp in raphé neuron (from Milescu et al., 2010). Right, Peak current vs. command voltage (red; the black curve is obtained when axonal Na channels are inactivated with a prepulse).

**Figure 5.**
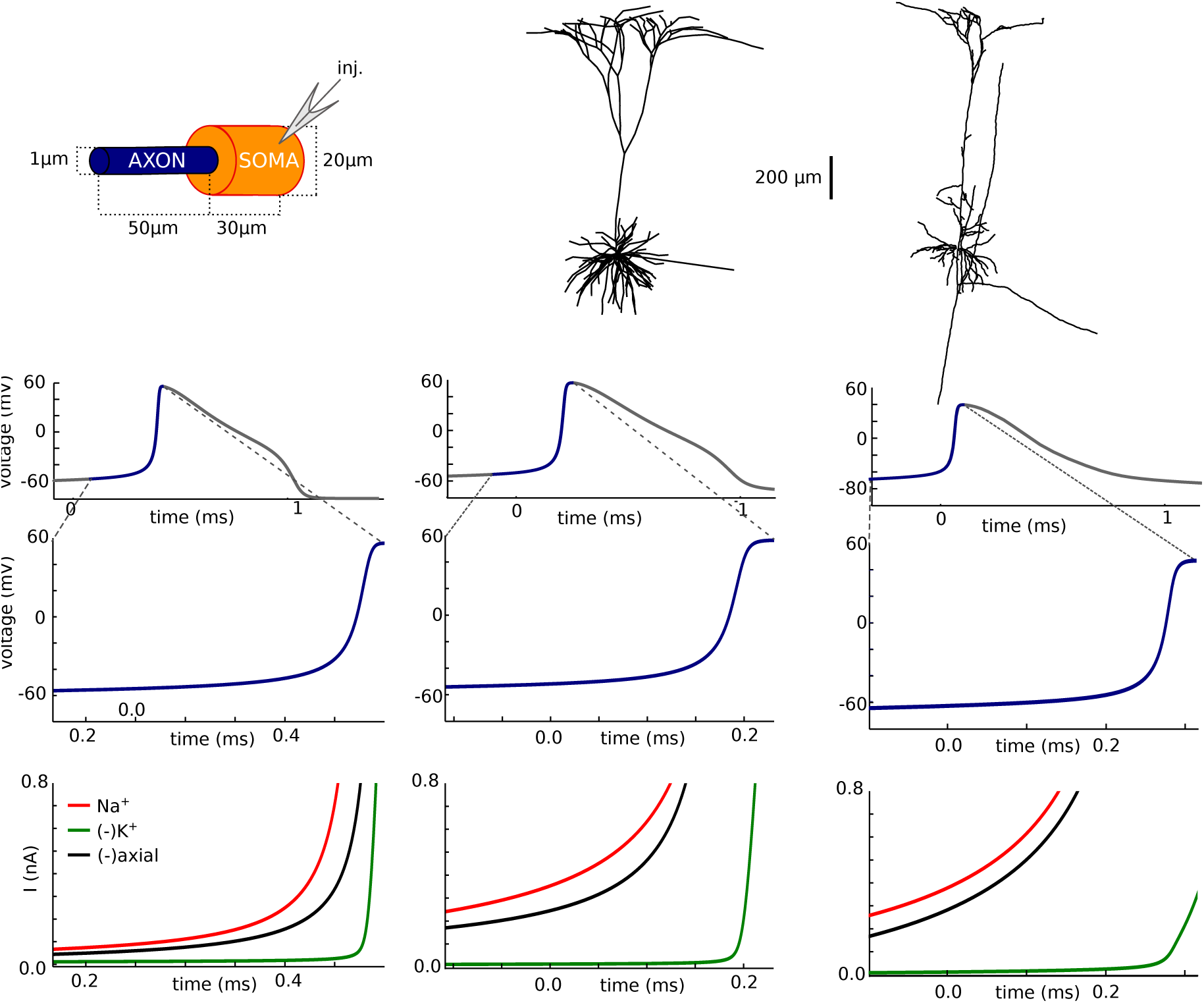
Balance of currents at spike initiation. A, Axonal spike. B, Rising phase of the spike. C, Na (red), K (green) and axial (green) current traces at the axonal initiation site. Na and K currents are summed over the AIS membrane; axial current is measured at the soma-axon junction.

### Excitability increases with intracellular resistivity

A consequence of this unconventional balance of currents is that the axial resistance between the soma and initiation site has a direct and possibly counter-intuitive impact on excitability: if axial resistance is increased, the neuron should become *more* excitable, despite the fact that the electrotonic distance of the initiation site increases. Axial resistance is proportional to the resistivity Ri of the intracellular medium. We therefore tested this prediction by manipulating the intracellular resistivity Ri in the model (Fig. 6A). When Ri is increased to 250 Ω.cm (orange) compared to 150 Ω.cm originally (green), spikes are initiated at a lower voltage. Conversely, when Ri is decreased to 30 Ω.cm, spikes are initiated at a higher voltage (light blue). When Ri is decreased further to 1 Ω.cm, the kink at spike onset disappears (dark blue). In all these cases except when R_i_ = 1 Ω.cm where the cell is essentially isopotential, spikes initiate in the axon.

**Figure 6.**
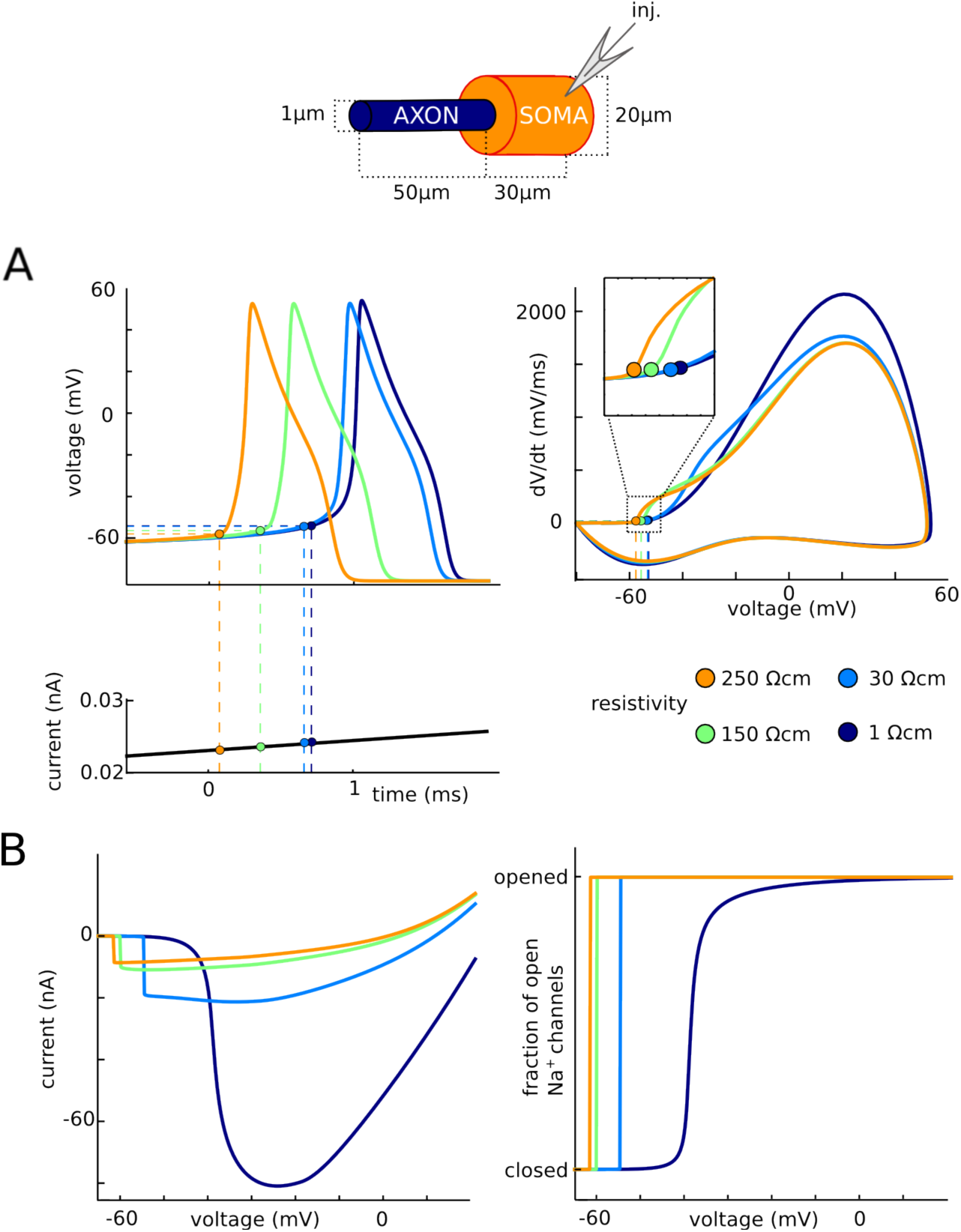
Impact of intracellular resistivity Ri on excitability. A, Spikes are triggered by a slow current ramp, for different values of Ri between 1 Ω.cm and 250 Ω.cm (green: original value). The neuron is more excitable for larger values of Ri. B, Current vs. somatic voltage in somatic voltage-clamp (as in Fig. 2B) and fraction of open Na channels vs. somatic voltage, for different values of Ri.

The same effect is seen in somatic voltage-clamp (Fig. 6B): the discontinuity in current is seen at increasingly higher voltages as resistivity decreases. These curves also demonstrate another feature of resistive coupling: as resistivity decreases, the peak current increases. This occurs because the resistive current is inversely proportional to resistance, by Ohm's law. Thus, as resistivity increases, the neuron becomes less excitable (higher threshold) but the axon transmits a larger current at initiation.

Finally, when intracellular resistivity is decreased further (Fig. 6B, dark blue), the phenomenon disappears, as predicted by the critical resistive coupling hypothesis: axonal current and proportion of open Na channels vary gradually with somatic voltage. In summary, excitability depends on the resistance between soma and AIS, and not only on local membrane properties of the initiation site.

### Sharp spike initiation requires a large enough somatodendritic compartment

A requirement of critical resistive coupling is that the soma effectively clamps the voltage at the beginning of the axon at spike initiation, which can occur if the somatodendritic compartment is large enough (Brette, 2013). We therefore show how spike initiation is affected by changing soma size in the simple model (Fig. 7). As was previously noted (Yu et al., 2008), the kink at spike onset entirely disappears when the soma has the same diameter as the axon (Fig. 7A, left; phase slope at 20 mV/ms: 5 ms^-1^) and only appears when the soma is large enough (Fig. 7A, middle and right; phase slope at 20 mV/ms: 12 and 31 ms^-1^; maximum phase slope of first component: 50 and 42 ms^-1^), or when a dendrite is added (Yu et al., 2008). Yet in all cases, when the soma is voltage-clamped, both the Na current and the proportion of open Na channels change abruptly when the somatic voltage exceeds a threshold (Fig. 47). This phenomenon is not explained by backpropagation.

**Figure 7.**
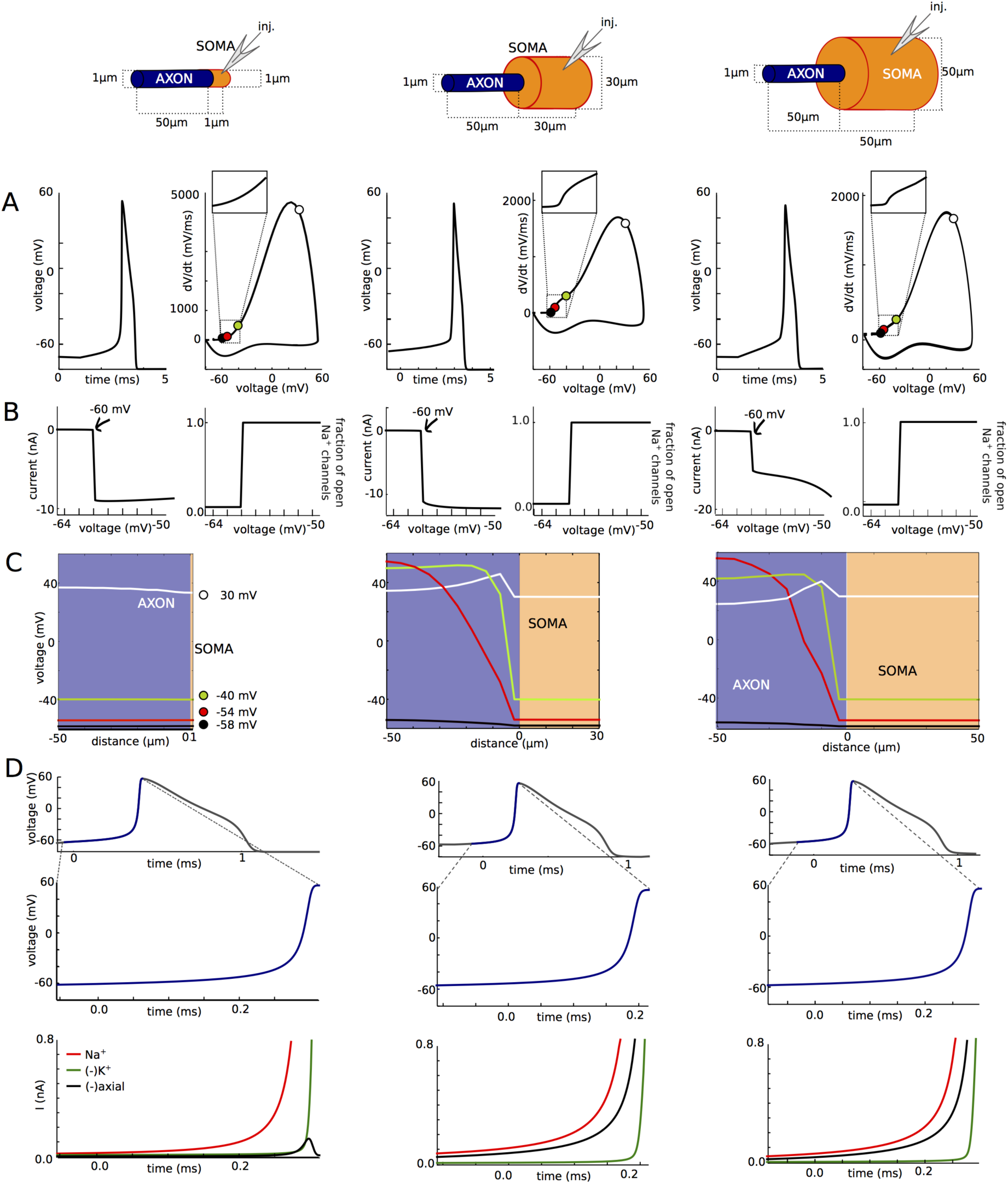
Influence of soma size on spike initiation. A, Somatic voltage trace and phase plot. B, Peak Na current (left) and proportion of open axonal Na channels (right) versus holding potential in somatic voltage-clamp. C, Membrane potential across the neuron at different instants near spike initiation. D, Balance of currents at initiation site (bottom) near spike initiation (top and middle: voltage trace).

Why does the discontinuity in somatic voltage-clamp result in sharp spike onsets when the soma is large but not when it is small? On Fig. 7C, we show the voltage along the axon at different moments of the spike upstroke, when somatic voltage is -58, -54, -40 and 30 mV (corresponding to the colored disks on Fig. 7A). In the case of the uniform axon (left), the neuron is essentially isopotential: the entire axon is depolarized synchronously. The situation is different when the soma is large (middle and right): at spike initiation, the soma almost clamps the proximal end of the axon while the voltage at the distal end rises with the Na influx. That is, the somatic currentclamp configuration corresponds, from the viewpoint of the axon, to a voltage-clamp of the start of the axon at the time scale of spike initiation. Thus the current discontinuity seen in somatic voltage-clamp (Fig. 7B) also appears in current-clamp near spike initiation, resulting in the sharp onset of spikes (Fig. 7A). Accordingly, the axial current is the main current opposing the Na current at the initiation site, meaning that the soma is a current sink for the initiation site, which is not the case with the uniform axon, where the axial current is small (Fig. 7D).

### Backpropagation does not sharpen spikes

Next, we show that the spike does not actually sharpen as it travels to the soma, and furthermore onset rapidness does not depend on somatodendritic capacitance, once the basic phenomenon is present. The sharp somatic spike onset is not a sharpened axonal spike onset, but rather reflects the maximum rapidness of the axonal spike, observed at a higher voltage.

We first make a methodological point. The standard way of measuring onset rapidness is to calculate the slope of the phase plot (dV/dt vs. V) at an arbitrary value of dV/dt (typically 5-20 mV/ms). In real somatic recordings, the phase plot is approximately linear over a wide enough range of dV/dt values, so that the exact choice is not critical (Baranauskas et al., 2010) (see Fig. 2F therein), all the more than it generally corresponds to only a few data points. However, in models where morphological parameters are varied over several orders of magnitude, this choice of dV/dt can be important. Fig. 8A (left) shows the phase plot of a spike in the simple model, for a somatic area of 3,000 μm^2^ (grey) and 10,000 μm^2^ (orange). It appears that the phase plot is linear around different values of dV/dt in the two cases (25 mV/ms and 60 mV/ms). When measured in the linear region of the phase plot, phase slope is similar in the two cases (40 and 50 ms^-1^). But when measured at the same value of dV/dt, phase slope can be very different in the two cases: at 20 mV/ms, it is about 3 times larger with the larger soma than with the smaller soma (Fig. 8A, right). This is artifactual because the measurements are done in different parts of the spike. Therefore, we defined onset rapidness as the phase slope in the linear part of the phase plot, which corresponds to the maximum phase slope of the first component of the phase plot.

**Figure 8.**
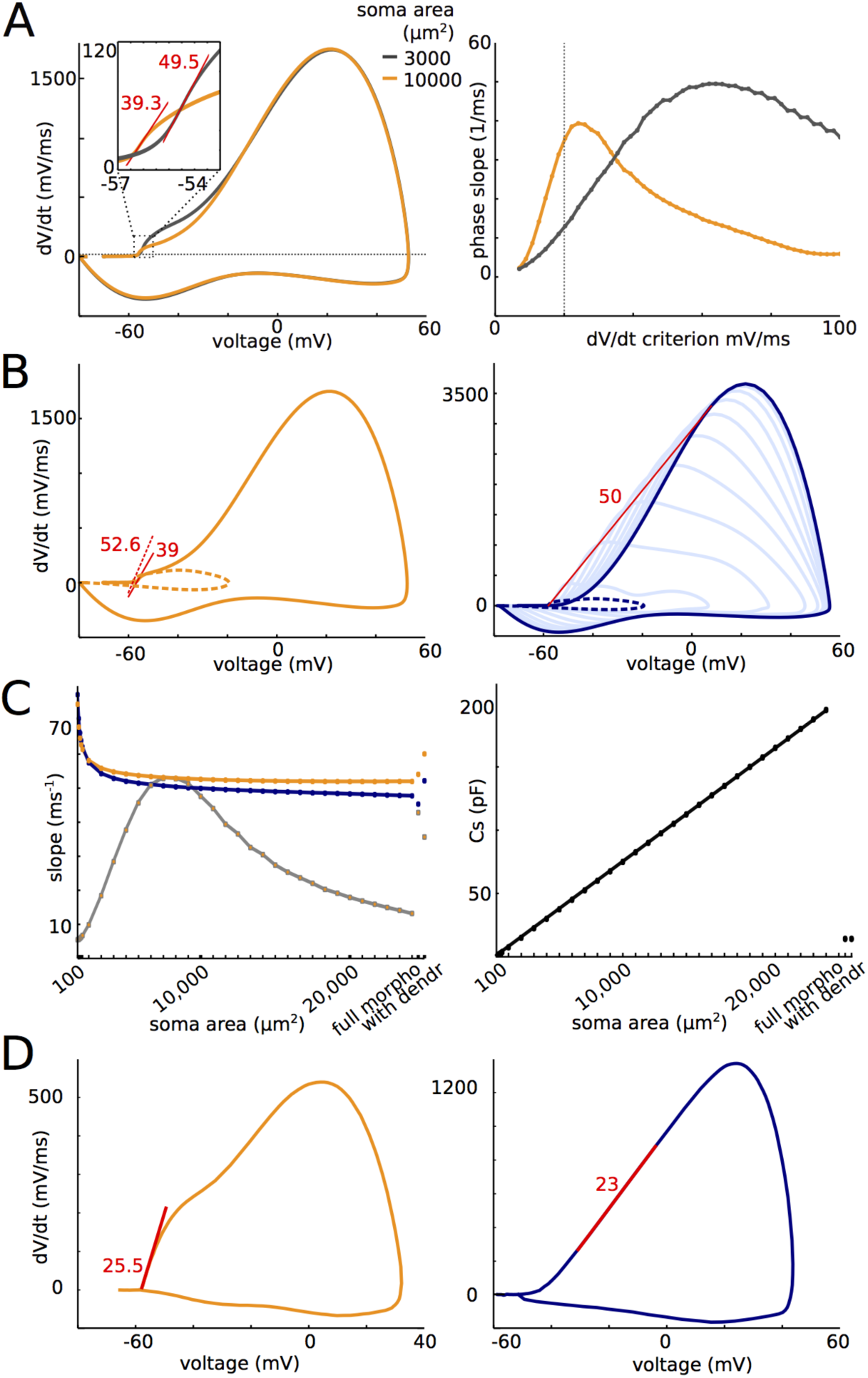
Somatic onset rapidness. A, Phase plot of a somatic spike with a large soma (orange, 10,0 μm^2^) and a small soma (gray, 3,000 μm^2^). The phase plot is linear (corresponding to locally constant phase slope) around dV/dt = 25 mV/ms in the former case and around 60 mV/ms in the latter case. Maximum phase slope is similar in both cases (39.3 and 49.5 ms^-1^). B, Left: phase plot for a large soma (10,000 μm^2^). The presence of somatic Na channels slightly decreases onset rapidness (orange, slope: 39 ms·^1^). Without them, onset rapidness is 52.6 ms·^1^. Right: phase plots at different points along the axon (dotted blue: soma; dark blue: distal end; light blue: intermediate axonal positions). The prediction of somatic onset rapidness based on resistive coupling is the maximum slope of a tangent to the phase plot intersecting the spike initiation point, which gives 50 ms^-1^ at the distal end (red line). C, Left: Somatic onset rapidness (orange) and prediction from axonal phase plots (blue) as a function of soma area for the simple model. The morphologically detailed model and the simple model with a dendrite are also shown on the right. Grey: somatic phase slope at 20 mV/ms. Right: For comparison, total somatic capacitance is shown as a function of soma area. D, Somatic (left) and axonal (right) phase plots of a spike digitized from Yu et al. (2008). Maximum phase slopes are similar.

To isolate the contribution to onset rapidness due to the axonal current, in the following analysis we removed somatic Na channels from the models. As is shown on Fig. 8B for a somatic area of 10,0 μm^2^ (left), the presence of somatic Na channels makes a small but significant difference in onset rapidness (39 ms^-1^ with and 53 ms^-1^ without). Fig. 8B (right) shows the axonal spike at the distal initiation site (dark blue) and at different places along the axon (light blue), when somatic Na channels are removed. It appears that the maximum dV/dt decreases approximately linearly as it travels to the soma, which can be directly explained by a resistive effect. In the critical resistive coupling hypothesis, the soma is driven at spike initiation by an axonal current that is essentially resistive, so that somatic onset rapidness should be determined by properties of the axonal spike at the initiation site. Specifically, somatic onset rapidness should equal the slope of a tangent to the axonal phase plot passing through spike threshold (see Methods), a value of the same magnitude as the maximum phase slope. As is shown on Fig. 8B (red), this theoretical prediction is satisfied in this model (50 ms^-1^ vs. 53 ms^-1^). In addition, the theory predicts that the same should hold at all axonal points between initiation site and soma, which is also approximately the case here (the red line is also almost tangent to all light blue curves). Thus, spikes do not sharpen as they travel to the soma; rather, the maximum phase slope is reached at lower and lower voltages.

This theoretical prediction matches somatic onset rapidness when somatic area is varied over several orders of magnitude (Fig. 8C, left). In fact, it can be seen that, as soon as somatic area is larger than a few hundred μm^2^, somatic onset rapidness (orange) does not depend much on somatic area. For comparison, Fig. 8C (right) shows the change in total somatic capacitance over the same variation in somatic area. It may appear so only when onset rapidness is measured at a fixed value of dV/dt (grey), for the reasons explained above, as also observed by Eyal et al. (2014) in a model with a dendrite of varying size. Note that the high values of onset rapidness for small soma are somewhat artifactual because they correspond to a case when the somatic phase plot is monophasic and maximum onset rapidness is reached at very high voltages (Fig. 7, left column).

We examined the simultaneous recordings of a spike in the soma and AIS bleb shown in (Yu et al. 2008) so as to test the theoretical prediction. Fig. 8D shows the digitized phase plots of the spike measured at the two sites. In the soma (orange), onset rapidness was about 25 ms^-1^. In the AIS (blue), the phase plot was nearly linear in the rising phase, with a slope of 23 ms^-1^ (note the different vertical scale). This match supports the resistive coupling hypothesis.

Thus, neither active backpropagation nor capacitive effects of the somatodendritic compartment sharpen spikes. Rather, the same value of maximum rapidness is reached at a lower voltage in the soma than in the axon. In fact, not only are spikes not sharpened by propagation, but their maximum slope (dV/dt) is in fact scaled down as they approach the soma (Fig. 8B, right), in agreement with the resistive coupling theory and with direct axonal measurements (Kole et al. 2008; Fig. 5b therein).

### Active backpropagation is not necessary for sharp spike initiation

We have seen that active backpropagation is not sufficient for sharp spike initiation. We next show that it is also not necessary (Fig. 9), both in the simple model (left column) and the morphologically detailed model (right column). We move all axonal Na channels to the same compartment, thereby suppressing active backpropagation. The exact result depends on the location of that compartment, in agreement with theoretical predictions (Brette, 2013), but in all cases, phase plots are biphasic, with initial onset rapidness between 46 and 71 ms^-1^ for the simple model and between 56 and 76 ms^-1^ for the detailed model (Fig. 9A). In detail, the voltage at spike onset decreases with increasing distance of the Na channels, and so does the maximum dV/dt in the first component of the phase plot. These features appear more clearly in somatic voltage-clamp (Fig. 9B). In all cases, Na channels open as a step function of somatic voltage. The spike threshold, corresponding to the discontinuity point in the current-voltage relationship, decreases when Na channels are placed further away, and peak axonal Na current also decreases with increasing distance. Thus the neuron is more excitable when Na channels are placed further away, but the axonal current transmitted to the soma is smaller. These two features are predicted by critical resistive coupling because the resistive axo-somatic current is smaller when Na channels are further away, so that a smaller Na current is required to trigger a spike, and a smaller resistive current is transmitted to the soma (Brette 2013). Despite these quantitative variations with the distance of the initiation site, the sharpness of spike initiation is unaffected by the exact distance: except when Na channels are very close to the soma, somatic onset rapidness is essentially independent of Na channel distance, around 70 ms^-1^ (Fig. 9C). This value is also close (in fact slightly higher) to the value obtained when Na channels are distributed across the AIS (about 50 ms^-1^). Therefore active backpropagation is not necessary to produce sharp spikes, and in fact does not contribute in making spikes sharper.

**Figure 9.**
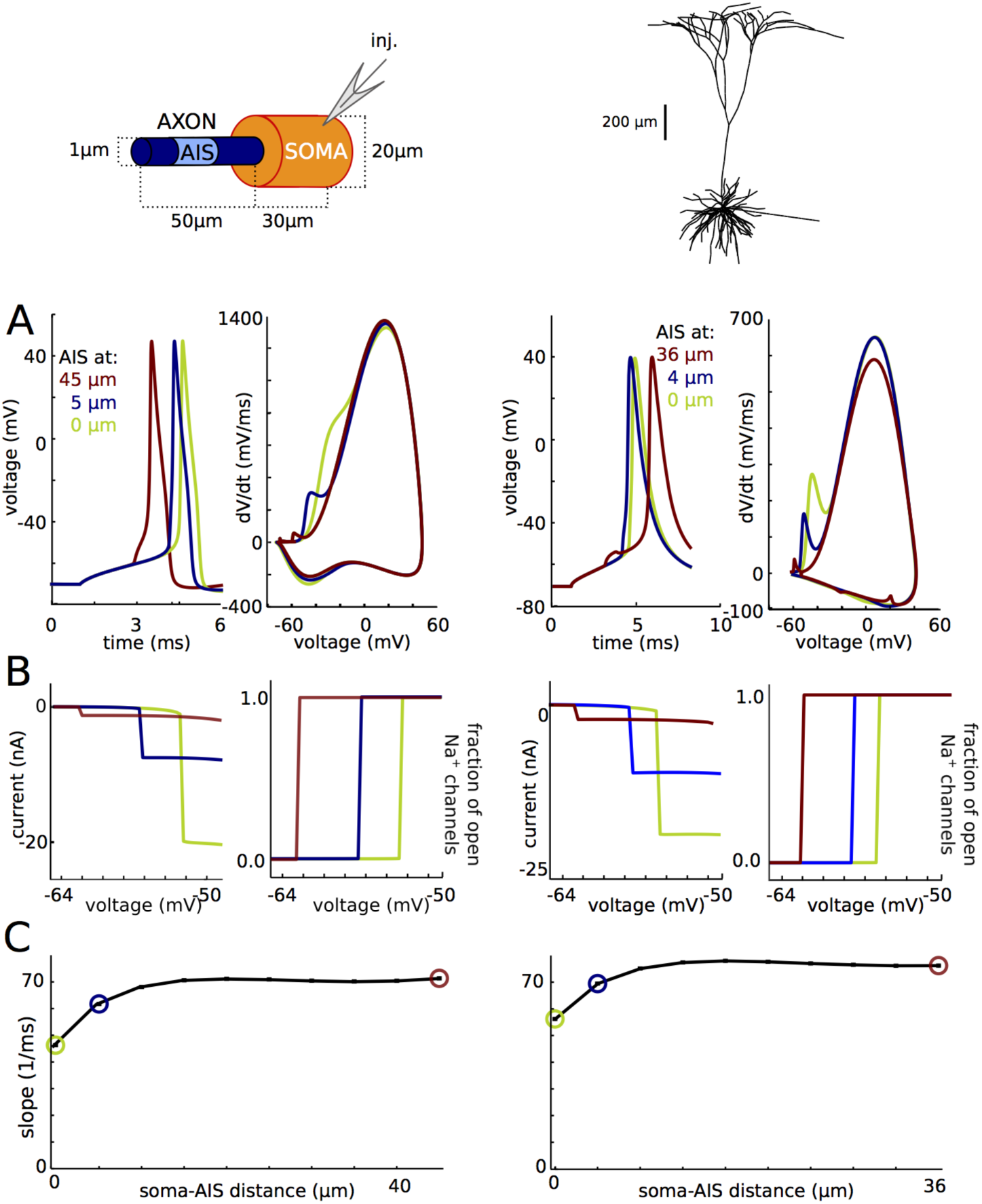
Active backpropagation is not necessary for sharp initiation. Left: model with simplified soma-axon geometry. Right: cortical pyramidal cell model with morphological reconstruction. A, Axonal Na channels are moved to a single axonal location (3 different locations shown). Left: voltage traces; right: phase plots. B, Peak Na current (left) and proportion of open axonal Na channels (right) versus holding potential in somatic voltage-clamp. C, Onset rapidness as a function of AIS position.

### Sharp somatic onset is reproduced by a model with two resistively coupled compartments

Finally, we designed a minimal two-compartment model that displays these features (Fig. 10A). The model included only Na and K channels with voluntarily highly simplified kinetics, with single gates and voltage-independent time constants, so as to show that the phenomenon is due to critical resistive coupling and not to specificities of channel kinetics (equilibrium values of gating variables shown in Fig. 10A). The maximum somatic Na conductance determines the threshold for spike regeneration in the soma (second component of the phase plots) (Platkiewicz and Brette, 2010); it was set at g_Na_^soma^ = 800 nS. For a somatic area of 2000 μm^2^ (corresponding to a 30 μm by 20 μm cylinder), this value corresponds to a conductance density of 400 pSμm^2^. The somatodendritic compartment is connected to the AIS compartment by a resistance R_a_. As shown in Fig. 9, this value determines the maximum dV/dt in the first component of the somatic phase plot (by Ohm's law, it is inversely proportional to Ra), and we chose R_a_ = 4.5 ΜΩ. This value corresponds to the onset of the AIS, close to the soma (rather than the distal initiation site), that is, the closest location where we expect to see a full spike at spike initiation. Finally, spike threshold is determined by the product g_Na_^axon^.Ra (Brette, 2013), and we chose g_Na_^axon^ = 1200 nS accordingly. For an AIS area of 50 μm^2^, this corresponds to a conductance density of about 7500 pS/μm^2^. Again, this value should be considered as an effective value and an overestimation of true conductance density, because distal channels have a greater impact on spike threshold and therefore require less conductance (Ra is greater at the distal end).

**Figure 10.**
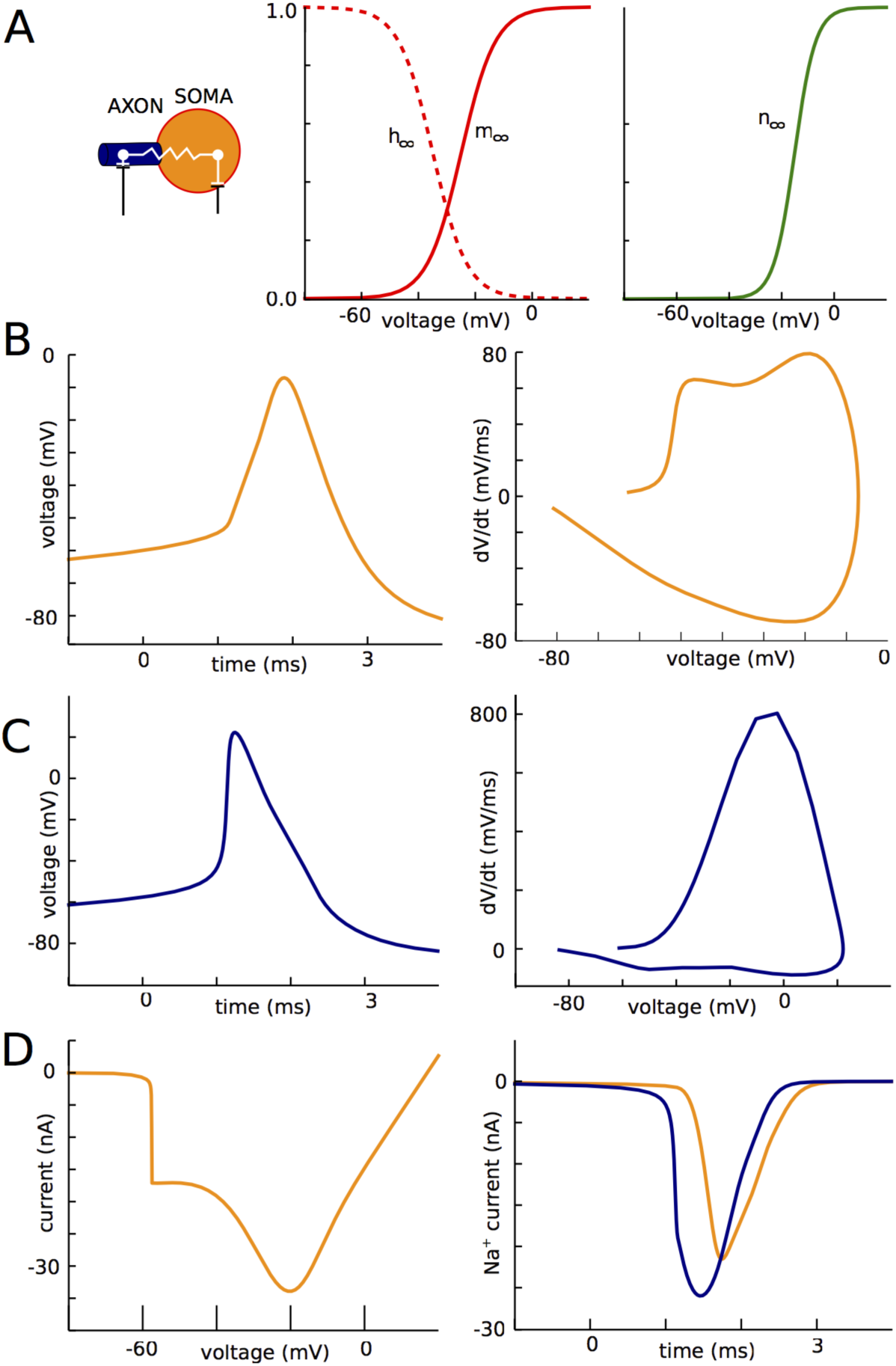
Two-compartment model. A, Equilibrium values of the gating variables for the Na (left) and K (right) channels. B, Voltage trace (left) and phase plot (right) of a somatic spike. C, Voltage trace (left) and phase plot (right) of an AIS spike. D, Left: Peak current recorded in somatic voltage-clamp as a function of holding voltage. Right: Na current in the AIS (blue) and soma (orange) during a spike in current-clamp.

With these parameter values, which are all within reasonable physiological ranges, we can see on Fig. 10B that the model exhibits sharp somatic spikes (onset rapidness: 17 ms^-1^) and a biphasic somatic phase plot, while in the AIS (Fig. 10C) spikes are smooth. The model also has a discontinuous current-voltage relationship measured in somatic voltage-clamp (Fig. 10D, left), as experimentally observed (Milescu et al., 2010). Finally, despite the order of magnitude difference between somatic and axonal Na conductance densities, total Na influxes in the AIS and in the soma are comparable (50% larger in AIS; Fig. 10D, right), as experimentally observed (Fleidervish et al., 2010). This occurs because the total conductances over each of the two compartments are in fact comparable. In detail, spike shape is not identical to measurements and this is expected given the simplicity of the model, but all features of sharp spike initiation are present even though the transmission of the axonal spike to the soma is purely resistive.

## DISCUSSION

By examining multicompartmental Hodgkin-Huxley models of spike initiation that reproduce the sharp onset of somatic spikes, we have found that active backpropagation and capacitive currents have no role in this phenomenon. Previous studies proposed that active channels between the initiation site and the soma increase spike onset rapidness. We have shown that entirely suppressing those currents has no effect on somatic onset rapidness. It has also been proposed that spike onset rapidness is further increased by the capacitance of the somatodendritic compartment (Yu et al., 2008; Eyal et al., 2014). We have shown that the increased somatic onset rapidness with larger capacitances observed in models is an artifact resulting from measuring phase slope at a fixed arbitrary value of dV/dt. When defined as the phase slope in the linear part of the phase plot, somatic onset rapidness does not change when capacitance is increased beyond a critical value. Finally, somatic onset rapidness can be directly predicted from spike shape at the distal initiation site (as maximum phase slope). Therefore, the sharpness of somatic spike onset in these models does not result from the properties of spike propagation, but rather of spike initiation.

Our analysis supports a biophysical explanation based on resistive coupling between the soma and the initiation site, where the soma effectively clamps the voltage at the start of the axon at spike initiation. According to this account, and as we have shown in these models, the main current opposing the Na current at spike initiation is not the transmembrane K current (i.e., mostly the leak), but the axial resistive current between the soma and initiation site. As a result, spikes initiate by the soma and AIS forming an electrical dipole, with current flowing between the two poles and charging the soma. Therefore, the phenomenon is not well modeled by the propagation of an electrical wave, since its wavelength would be the entire system. The observed delay between axonal and somatic spikes is thus better understood as a charging time than a propagation delay. One implication is that spike initiation is *not* a local axonal event, and therefore its characteristics are determined by the properties of the soma-AIS system, in particular its geometry. Specifically, the sharpness of spike initiation arises from the geometrical heterogeneity of that system. In a simplified two-compartment model representing the soma-AIS dipole, it was previously shown that spikes are initiated abruptly if the product of axial resistance and maximum Na conductance exceeds a critical value (Brette, 2013). Phenomenologically (but not biophysically), the corresponding model is mathematically quasiequivalent to the cooperative gating model of spike initiation (Naundorf et al., 2006) and therefore shares all its functional properties. A major common feature between these two models, supported by our analysis of the biophysical models and by experiments (Milescu et al., 2010), is that peak axonal current varies abruptly with somatic voltage, while current latency is inversely related to the difference between somatic voltage and threshold. A specific feature of the resistive model, also observed in the biophysically detailed models, is that increasing axial resistance, and therefore reducing axial currents, makes the neuron more excitable. This feature has been previously suggested to underlie structural tuning of the AIS (Kuba, 2012).

As we have mentioned, an alternative hypothesis is that Na channels in the AIS cooperate, so that they effectively open all at once when the axonal voltage exceeds a threshold (Naundorf et al., 2006). Cooperative activation has been demonstrated in calcium (Marx et al., 1998, 2001), potassium (Molina et al., 2006) and HCN channels (Dekker and Yellen, 2006), and in pharmacologically altered Na channels of cardiac myocytes (Undrovinas et al., 1992). It should appear in AIS phase plots as a very large increase in dV/dt at spike initiation (Öz et al., 2015), with a biphasic phase plot if only part of the channels (e.g. the Nav1.6 subtype) cooperate (Huang et al., 2012). However, phase plots of spikes recorded in axonal blebs near the AIS are monophasic, with a gradual increase with voltage as expected with non-cooperative channels (McCormick et al., 2007; Yu et al., 2008). In isolated blebs, TTX has no effect on half-activation voltage of Na channels, whereas cooperativity predicts an increase (Hu et al., 2009). It has been opposed that cooperativity may depend on the intact cytoskeleton of the AIS, which might be disrupted in axonal blebs (Naundorf et al., 2007), but the axonal bleb recordings were performed simultaneously with somatic recordings, which did exhibit a distinct kink at spike onset. Finally, voltage traces recorded in whole cell patch clamp in intact axons appear very similar to axonal bleb recordings and show no sign of cooperativity, with a smooth spike onset at the AIS (Kole and Stuart, 2008). We conclude that at this date there is no evidence of cooperativity of Na channels in the AIS.

The cooperativity hypothesis was also motivated by the observation of input-output properties of cortical neurons that are not well accounted for by the standard account of spike initiation (Brette, 2015), in particular the fact that cortical neurons can transmit very high input frequencies (Ilin et al., 2013). The backpropagation account only addresses somatic spike shape, but not spike initiation *per se*. The critical resistive coupling model addresses both aspects because it is mathematically almost equivalent to the cooperativity model, although it has a different biophysical basis (axial resistance in the resistive model corresponds to channel coupling in the cooperativity model) and makes different predictions in the axonal initiation site.

We now discuss experimental evidence regarding the critical resistive coupling hypothesis, starting with the notion that the somatodendritic compartment is a current sink for the initiation site. First, the initiation site is very close to the soma, as previously noted, and axonal diameter is small compared to the soma, especially if the conductance load of the dendritic tree is considered (Hay et al., 2013). Thus, biophysical theory predicts that the soma is a current sink for the initiation site (Brette, 2013), i.e., that most Na current entering the AIS flows to the soma, producing a voltage gradient between the two sites. This prediction is backed up by several lines of evidence. First, there is generally no voltage gradient across the two sites between spikes, but this gradient rises to about 7 mV near spike initiation due to the opening of Na channels (Kole and Stuart, 2008), a value close to the theoretical prediction of ka (Boltzmann slope factor of Na channel activation) (Brette, 2013). Second, axonal outside-out patch-clamp recordings show that there is little K current flowing during the rising phase of the AIS spike (Hallermann et al., 2012). Third, Na imaging experiments show that the peak of Na influx during a spike occurs at an axonal position closer to the soma than the initiation site (Baranauskas et al., 2013). This is indeed expected if most Na current flows to the soma, because voltage then increases monotonously with distance from the soma (spatial gradient of voltage is proportional to axial current), so that spikes are initiated at the distal end of the AIS even though Na channel density may be lower.

Thus, there is convincing experimental evidence that the soma acts as a current sink for the initiation site, with most Na current flowing directly to the soma. Evidence that the product of axial resistance and Na conductance is large enough to produce an abrupt opening of Na channels is most directly provided by somatic voltage-clamp experiments, which show a discontinuity in the measured current-voltage relationship (Barrett and Crill, 1980; Milescu et al., 2010), although a finer resolution in the voltage commands would be desirable (voltage resolution is generally 5 mV). In practice, somatic voltage-clamp recordings are complicated by the fact that currents can be very large, and the precision of voltage clamping is limited to the product of current and uncompensated series resistance. Electrode artifacts could produce a discontinuity in observed current-voltage relationships when there is actually none. This was apparently not in the case in (Milescu et al., 2010) because continuous currents of the same magnitude were observed when axonal channels were inactivated with a prepulse. In that study, currents were reduced by applying a small dose of TTX. Indirect evidence about the discontinuous opening of Na channels, including somatic spike onset and input-output properties, is reviewed in (Brette, 2015). Finally, the comparison of somatic onset rapidness and axonal phase slope also matches theoretical predictions (Fig. 5D), although more experimental recordings would be required to test these predictions.

To test the critical resistive coupling theory more directly, one would need to manipulate the axial resistance between soma and initiation site. The neuron should be more excitable when axial resistance is increased, but the current transmitted to the soma should decrease. If the resistance were decreased very substantially, spike initiation should become smooth instead of sharp. We list here several experimental possibilities, chemical and mechanical, although none is particularly easy to perform. Intracellular resistivity could be changed by manipulating the ionic composition of the intracellular medium. This is difficult because it must be compensated by corresponding changes in the extracellular medium to preserve osmotic equilibrium, without disturbing reversal potentials, but the extracellular and intracellular media cannot be modified simultaneously. In a similar way, extracellular resistance could be manipulated: although typically neglected in models, it has the effect of increasing the total axial resistance. For example, Hodgkin demonstrated that conduction velocity decreased when the axon was immersed in oil, which reduces the water volume around the axon to a thin layer (Hodgkin, 1939). A similar experiment could be done to test changes in excitability. Another idea is to use osmotic pressure to change the diameter of the axon, thereby changing total axial resistance. However, it is possible that the dense cytoskeleton of the AIS provides rigidity so that changes in shape might be limited to the soma. Finally, pinching the base of the axon with two glass pipettes would increase axial resistance, and it has been done with dendrites (Bekkers and Haüsser, 2007). All of these experimental ideas are challenging, but not impossible in principle.

In conclusion, biophysical modeling and experimental evidence support the notion that, in neurons with an AIS, normal (orthodromic) spike initiation results from the interplay between the Na current and the resistive axial current flowing between the AIS and soma, rather than between local transmembrane Na and K (leak) currents as in somatic and ectopic initiation (in an axon far from the soma). This means that the mechanism of spike initiation is not local to the axon, but rather occurs through the formation of a resistively coupled soma-AIS dipole. This situation occurs because of the large variation in geometry and biophysical properties over a small spatial scale. Accordingly, at least in the biophysical models that we have examined, the kink at somatic spike onset results from strong coupling of the soma-AIS system, rather than from an artifact of somatic recording.

## MATERIALS AND METHODS

### Detailed neuron models

We used two spatially extended neuron models described in (Yu et al., 2008), a simple model of a cylindrical soma and a cylindrical 50 μm long axon (diameter 1 μm) available on ModelDB (Hines et al., 2004) and a morphologically detailed model based on a reconstruction of a cortical pyramidal cell (personal communication of Prof. Yuguo Yu). We also used a multicompartmental model of a pyramidal cell described in (Hallermann et al., 2012), also available on ModelDB. These models were implemented in Neuron 7.4 (Hines and Carnevale, 1997) with time step 1 μs and analyzed in Python. In Fig. 3, the extracellular field was computed with the standard line source method (Holt et al., 1999; Einevoll et al., 2013), with extracellular conductivity σ = 0.3 S.m^-1^, and simulated with the NeuronEAP Python library (Telenczuk et al., 2016).

#### Morphology

The simple model consists of an axon modeled as a cylinder of length 50 μm and diameter 1 μm, and a soma modeled as a cylinder of variable length and diameter (the bleb and dendrite were not included unless stated otherwise). In Fig. 2 and 6, the soma has diameter 20 μm and length 30 μm. In Fig. 4 and 5, soma size was varied with equal length and diameter. In one case (Fig. 4C–D), we kept the dendrite of length 3000 μm and diameter 5 μm (60 segments).

The first morphologically detailed model (Yu et al., 2008) is based on a reconstructed layer 5 cortical pyramidal cell. The axon consists of a 10 μm long axon hillock, with diameter tapering from 4.8 μm to 1.2 μm, connected to an initial segment of diameter 1.2 μm and length 40 μm, followed by a 500 μm myelinated axon with 5 nodes of Ranvier separated by 100 μm. The second morphologically detailed model (Hallermann et al., 2012) is also based on a reconstructed layer 5 cortical pyramidal cell, and we used the model as described in that reference.

#### Channel properties

Passive properties were set as in (Yu et al., 2008): specific membrane capacitance C_m_ = 0.75 pS/μm^2^, intracellular resistivity R_i_ = 150 H.cm, specific membrane resistance R_m_ = 30,000 Ω.cm^2^, leak reversal potential E_L_ = -70 mV, membrane time constant τ = 22.5 ms. In the simple model, Na conductance densities were 8000 pS/μm^2^ in the axon, 0 or 800 pS/μm^2^ in the soma and 20 pS/μm^2^ in the dendrite. In the morphologically detailed model, Na conductance densities were 8000 pS/μm^2^ in the hillock and in the axon initial segment, 800 pS/μm^2^ in the soma and 100 pS/μm^2^ in dendrites. In the simple model, we only included Na and potassium channels; in the morphologically detailed model, we used all the channels present in the original model. Detailed channel kinetics and properties of other channels are described in (Yu et al., 2008).

In Fig. 9 we moved all channels involved in spike generation (Na and potassium) to a single compartment while maintaining the same total conductance. As the initial segment consists of 10 compartments, this corresponds to multiplying conductance densities by 10 in the target compartment and setting them to 0 in all other compartments.

### Two-compartment model

The two-compartment model represents the soma and the axon initiation site, coupled by axial resistance R_a_ = 4.5 MH. Capacitances are C_s_ = 250 pF for the soma, in the range of values measured in layer 5 pyramidal neurons of rats (Arsiero et al., 2007), and C_a_ = 5 pF for the axon. The axonal value was chosen empirically, as in reality axon impedance is highly frequencydependent. Leak conductance is 12 nS (corresponding to a 20 ms membrane time constant) and leak reversal potential is -80 mV.

For ionic channels, we deliberately used the simplest possible models so as to show that the sharpness of spikes does not result from subtle aspects of their detailed properties (Fig. 10A). Na channel activation is modeled with first-order kinetics, half-activation -25 mV, Boltzmann slope 6 mV, time constant 100 μs (voltage-independent). Na channel inactivation is considered independent with time constant 0.5 ms, half-inactivation voltage -35 mV, Boltzmann slope 6 mV. K channels are also modeled with first-order kinetics, half-activation -15 mV, Boltzmann slope 4 mV, time constant 2 ms. Total Na conductance is 800 nS in the soma and 1200 nS in the axon; total K conductance is 2200 nS in the soma and 1200 nS in the axon.

The model was simulated with the Brian simulator 2.0 (Stimberg et al., 2014).

### Analysis

#### Voltage-clamp

In voltage-clamp measurements, the soma was clamped to a holding potential and the current was measured and corrected for the leak current with the P/n protocol, as in (Milescu et al. 2010; Benzanilla, 1977). The peak current is shown as a function of holding potential.

#### Phase slope

The standard way of measuring onset rapidness is to calculate the slope of the phase plot (dV/dt vs V) at a certain value of dV/dt (typically 5-20 mV/ms). In real somatic recordings, the phase plot is approximately linear over a wide enough range of dV/dt values, so that the exact choice is not critical (Baranauskas et al., 2010) (see Fig. 2F therein). However, in models where morphological parameters are varied over several orders of magnitude, the phase plot can be linear around different values of dV/dt (Fig. 8). Therefore, we defined onset rapidness as the phase slope in the linear part of the phase plot, which corresponds to the maximum phase slope. When the spike is regenerated at the soma (somatic Na channels), there are two local maxima and we choose the smaller one (closer to spike onset).

### Theoretical prediction of onset rapidness

From the resistive coupling hypothesis, we can derive a theoretical prediction about somatic onset rapidness. We first assume that the major somatic current at spike initiation is the axonal current, so that the membrane equation reads:

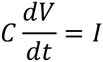

where C is membrane capacitance. Phase slope is then:

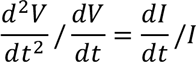

The resistive coupling hypothesis postulates that the axonal current is a resistive current:

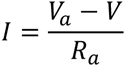

where V_a_ is axonal voltage and R_a_ is axial resistance between the two sites. It follows that:

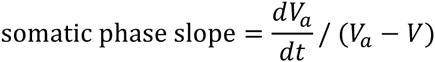

Assuming further that the axonal spike develops before the somatic spike, we consider that V is close to spike threshold. Onset rapidness is defined as the maximum phase slope (for the first component), as discussed above, and therefore should correspond to the maximum value of the formula above. Graphically, this maximum corresponds to the slope of a tangent to the axonal phase slope, starting from threshold (Fig. 8B, right). This value is in fact close to the maximum axonal phase slope. A simplified theoretical prediction is thus that somatic onset rapidness (or “initial phase slope”) approximately equals maximum axonal phase slope.

## Acknowledgments

We thank Yuguo Yu for sharing the code of his morphologically detailed model.

